# Propofol inhibits endogenous formyl peptide-induced neutrophil activation and alleviates lung injury

**DOI:** 10.1101/340711

**Authors:** Chun-Yu Chen, Yung-Fong Tsai, Wei-Ju Huang, Shih-Hsin Chang, Tsong-Long Hwang

## Abstract

Critically ill patients have a high risk of sepsis. Various studies have demonstrated that propofol has anti-inflammatory effects that may benefit critically ill patients who require anesthesia. However, the mechanism and therapeutic effect remain incompletely understood. Our previous data suggest that propofol can act as a formyl peptide receptor 1 (FPR1) antagonist. Here, we hypothesize that propofol mitigates sepsis-induced acute lung injury (ALI) by inhibiting mitochondria-derived *N*-formyl peptide-mediated neutrophil activation. In human neutrophils, propofol competitively reduced the release of elastase, superoxide, and reactive oxygen species induced by fMMYALF, a human mitochondria-derived *N*-formyl peptide. In addition, propofol significantly inhibited fMMYALF-induced chemotaxis, calcium mobilization, and phosphorylation of protein kinase B and mitogen-activated protein kinases. These results indicate that propofol suppresses neutrophil activation by blocking the interaction between endogenous *N*-formyl peptide and its receptor, FPR1, thus inhibiting downstream signaling. Furthermore, propofol alleviated alveolar wall disruption, edematous changes, and neutrophil infiltration in lipopolysaccharide-induced ALI in mice. Noticeably, propofol improved the survival of sepsis mice. This study indicates that the anti-neutrophil effects of propofol may benefit critically ill septic patients.

## Introduction

Propofol (2,6-Diisopropylphenol) is a commonly used anesthetic drug that is administered intravenously for the induction and maintenance of anesthesia. The favorable pharmacokinetic characteristics of propofol make it a rapid-onset and short-acting agent. In addition, the anti-inflammatory and antioxidant effects of propofol (Fan et al, 2015; Marik, 2005; Vanlersberghe & Camu, 2008) increase the advantage of its use in clinical practice. Several studies have indicated that propofol can moderate many aspects of inflammatory responses. Propofol suppresses the immune activities of monocytes/macrophages (Chen et al, 2003; Li et al, 2010; Marik, 2005; Tang et al, 2011; Vanlersberghe & Camu, 2008; Wheeler et al, 2011) and neutrophils (Galley et al, 1998; Marik, 2005; Mikawa et al, 1998; Yang et al, 2013), including chemotaxis, extravasation, migration, phagocytosis, and production of reactive oxygen species (ROS). Moreover, propofol attenuates proinflammatory cytokine generation (Liu et al, 2016; Marik, 2005; Taniguchi et al, 2002; Vanlersberghe & Camu, 2008) and reduces the biosynthesis of nitric oxide (Chen et al, 2003; Fan et al, 2016; Marik, 2005; Vanlersberghe & Camu, 2008; Xu et al, 2017; Yagmurdur et al, 2017) both *in vitro* and *in vivo*.

The risk of systemic inflammatory response syndrome (SIRS) and infection is higher among critically ill patients who have experienced trauma or cardiac arrest or have undergone surgery. Endogenous damage-associated molecular patterns (DAMPs), which are released in large amounts from damaged cells or tissues in critically ill patients, activate the innate immune system, and make a contribution to the pathogenesis of septic shock, acute lung injury (ALI), and multi-organ failure. Moreover, once the immune system cannot restrain an invading pathogen sufficiently, the overwhelming inflammatory responses may further deteriorate the immune system’s antimicrobial function, engendering a vicious cycle. Recently published protocols for managing critically ill patients proposed that continuous or intermittent sedation should be minimized (Rhodes et al, 2017). However, treating some critically ill patients, such as those who suffer from hypersensitive airway or increased intracranial pressure, requires the use of sedatives for providing bronchodilation (Calzetta et al, 2015; Grim et al, 2012) and neuroprotective effects (De Cosmo et al, 2005; Ulbrich et al, 2016) as well as minimizing stress responses. Moreover, anesthesia is an essential measure for inducing unconsciousness and analgesia in critically ill patients who required surgery. In summary, sedative agents should be selected cautiously for anesthetizing patients who are severely ill and at high risk of secondary infection and sepsis. The antioxidant and anti-inflammatory properties of propofol may benefit critically ill patients with SIRS.

Our previous study demonstrated that propofol is a competitive inhibitor of formyl methionyl-leucyl phenylalamine (fMLF) that functions by blocking formyl peptide receptor 1 (FPR1) (Yang et al, 2013). In this study, we hypothesized that propofol has a therapeutic effect through competitive inhibition of human neutrophil activation induced by an endogenous *N*-formylated peptides. Mitochondria-derived *N*-formylated peptides are quickly released following tissue or cell damage (Dorward et al, 2017; Rabiet et al, 2005; Raoof et al, 2010). These *N*-formylated peptides are strong chemoattractants and can initiate and aggravate inflammation, resulting in SIRS; thus, they can be considered as DAMPs. We executed the current study with the aim of ascertaining whether propofol inhibits fMMYALF, a human mitochondria-derived *N*-formylated peptide (Dorward et al, 2017; Rabiet et al, 2005), induced neutrophil activities, including respiratory burst, degranulation, and chemotaxis. Additionally, the pharmacological effects of fMMYALF were analyzed to evaluate whether the inhibitory effects of propofol are attributable to blocking of the interaction between fMMYALF and its receptor, FPR1, which interrupts receptor-mediated downstream signaling. Sepsis is the most common cause of ALI (Fowler et al, 1983), which results in increased lung permeability, enhanced neutrophil recruitment, respiratory failure, and mortality. We further investigated the protective effects of propofol in a murine model of ALI induced by endotoxin.

## Results

### Propofol competitively inhibits fMMYALF-induced elastase and superoxide release in human neutrophils

To assess whether propofol affects neutrophil function and inflammatory activities induced by endogenous mitochondrial-derived formyl peptide, we fist evaluated its effect on elastase and superoxide release in fMMYALF-activated human neutrophils. Our results revealed that propofol (5–100 μM) dose-dependently reduced elastase release and superoxide release with IC_50_ values of 6.00 ± 1.02 μM and 15.09 ± 3.65 μM, respectively (Fig 1A and 2A). Furthermore, the addition of propofol caused a parallel right shift in the dose–response curves of fMMYALF for elastase and superoxide release (Fig 1B and 2B), suggesting that propofol has a competitive effect with endogenous mitochondrial-derived formyl peptide.

**Figure 1.**
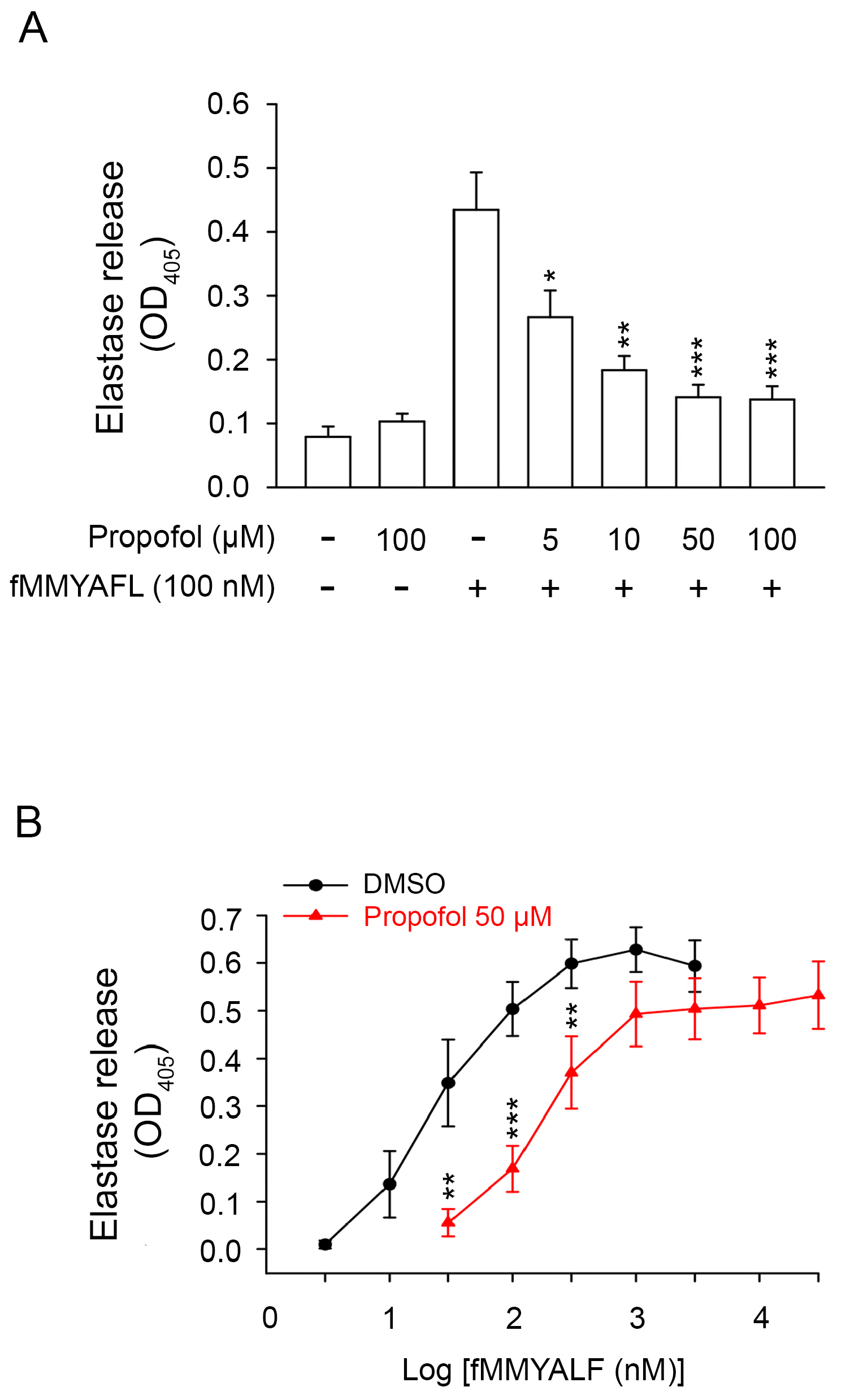
Propofol competitively inhibits fMMYALF-induced elastase release in human neutrophils. A, B Elastase levels in fMMYALF-induced human neutrophils. (A) concentration effects of propofol (n = 5). (B) dose–response curves of fMMYALF (n = 7). Data information: Data are expressed as the mean ± SEM. *p< 0.05, **p<0.01, ***p< 0.001 (Student’s t-test).

**Figure 2.**
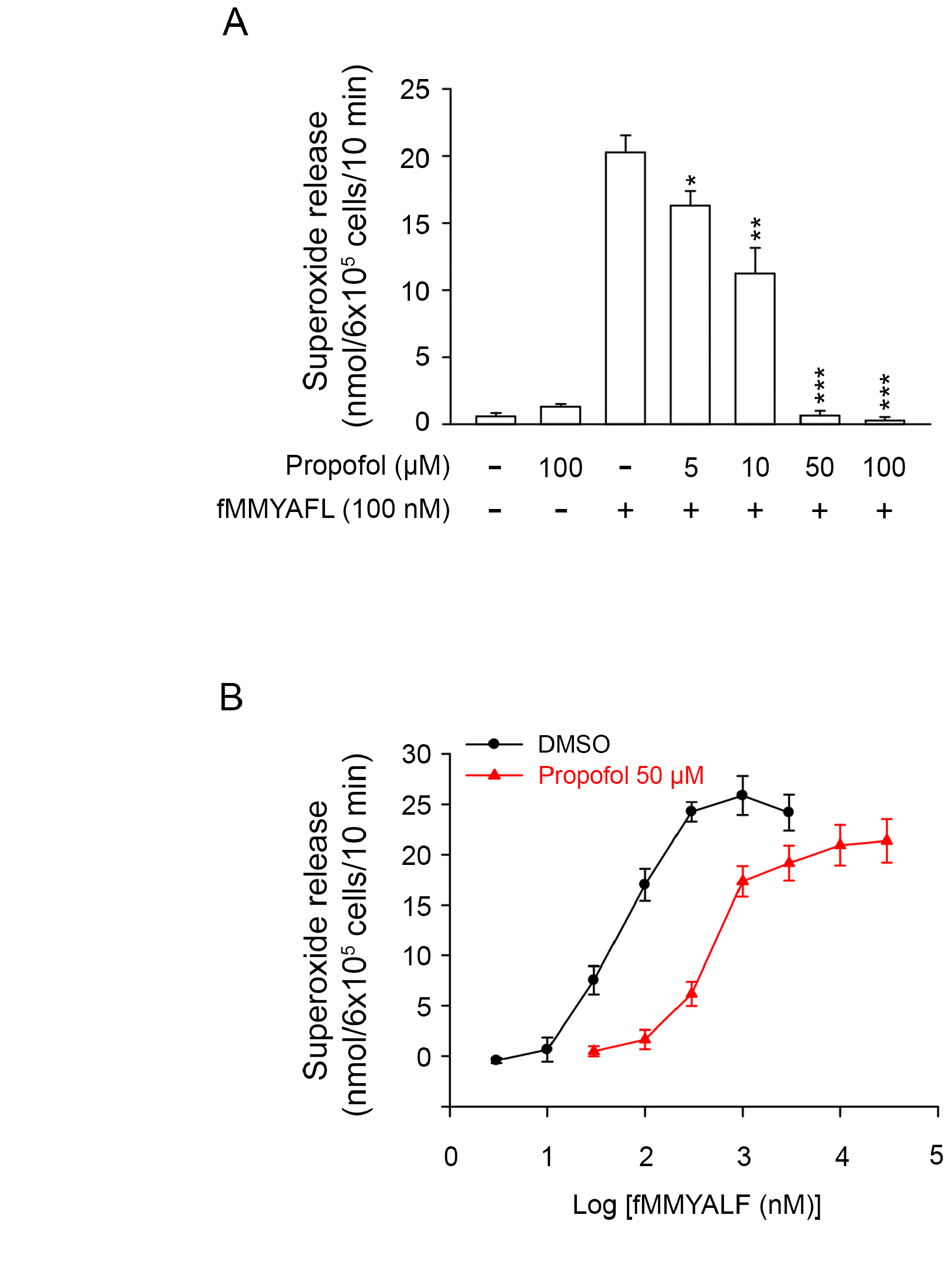
Propofol competitively inhibits fMMYALF-induced superoxide release in human neutrophils. A, B Superoxide release levels in fMMYALF-induced human neutrophils. (A) concentration effects of propofol (n = 6). (B) dose–response curves of fMMYALF (n = 6). Data information: Data are expressed as the mean ± SEM. **p<0.01, ***p< 0.001 (Student’s t-test).

### Propofol competitively reduces inhibited fMMYALF-induced ROS production and release in human neutrophils

In human neutrophils, the NADPH-oxidase-produced superoxide anion can be converted to many ROS that have strongly germicidal effects but are concomitantly harmful to tissues. Our results revealed that intracellular ROS production in fMMYALF-activated neutrophils was inhibited by propofol (50 μM). Propofol (50 μM) did not affect the basal ROS production in unstimulated neutrophils (Fig 3). Moreover, our data showed that propofol reduced the extracellular level of total ROS, which was measured using luminol-enhanced chemiluminescence method (Fig 4). Furthermore, the mixture of fMMYALF and propofol (50 μM) did not increase the release of lactate dehydrogenase (LDH) (data not shown), which suggests that the combined use of fMMYALF and propofol was not cytotoxic to human neutrophils.

**Figure 3.**
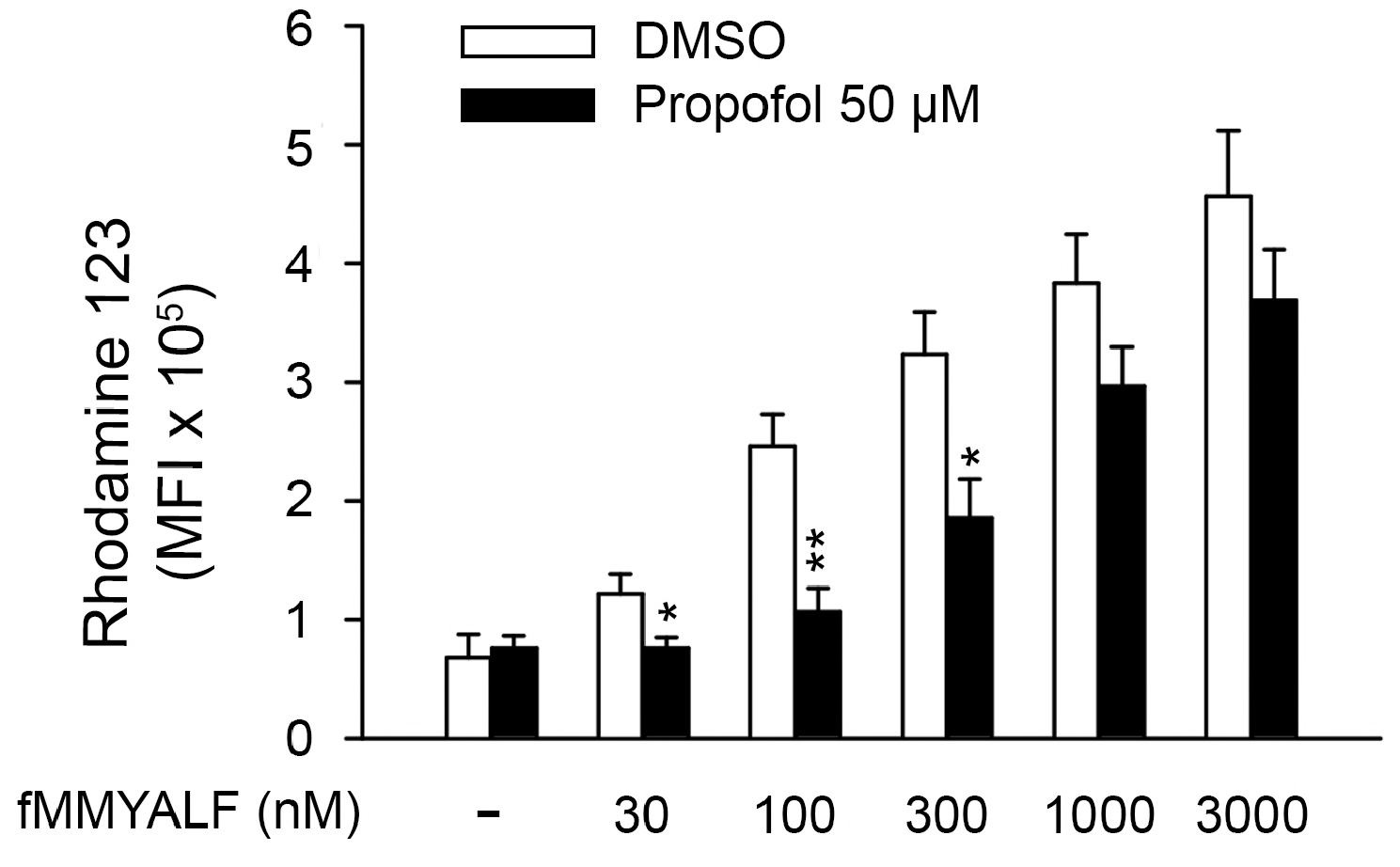
Propofol competitively inhibits fMMYALF-induced intracellular ROS production in human neutrophils. Intracellular ROS concentration in fMMYALF-induced human neutrophils (n = 6). Data information: Data are expressed as the mean ± SEM. *p< 0.05, **p<0.01 (Student’s t-test).

**Figure 4.**
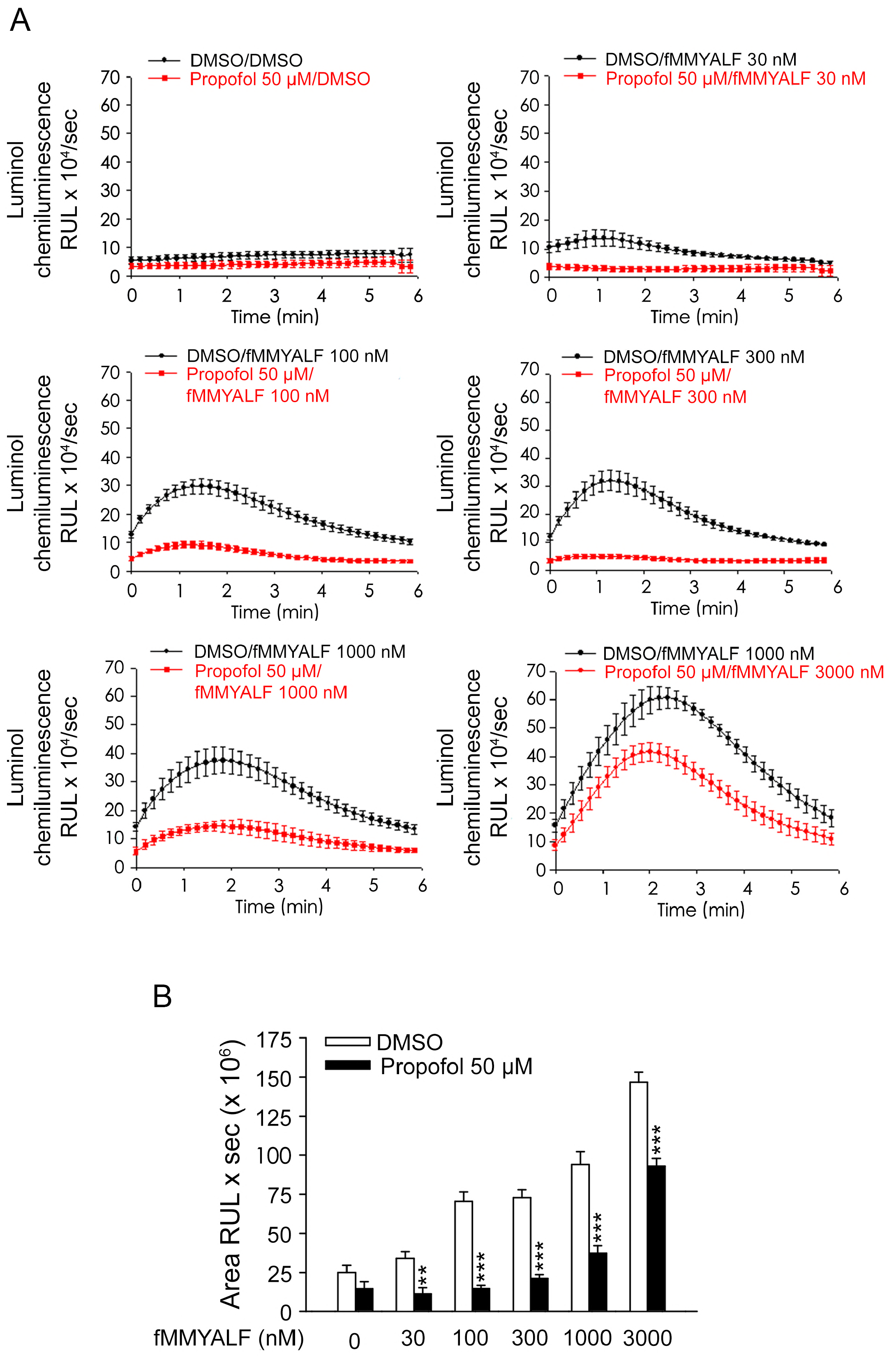
Propofol reduces total ROS release in fMMYALF-induced human neutrophils. A, B Total ROS levels in fMMYALF-induced human neutrophils. (A) Representative images of different concentrations of fMMYALF (n = 4 for each group). (B) Quantification of the levels of (A). Data information: Data are expressed as the mean ± SEM. **p<0.01, ***p< 0.001 (Student’s t-test).

### Propofol inhibits chemotaxis of human neutrophils

The extravasation and migration of neutrophils into inflammatory sites is a critical response in FPR1-mediated immunological events. For investigating whether propofol reduces neutrophil migration, we examined the chemotactic response of neutrophils to fMMYALF. Our results demonstrated that propofol (50 μM) significantly reduced the fMMYALF-stimulated migration of neutrophil (Fig 5).

**Figure 5.**
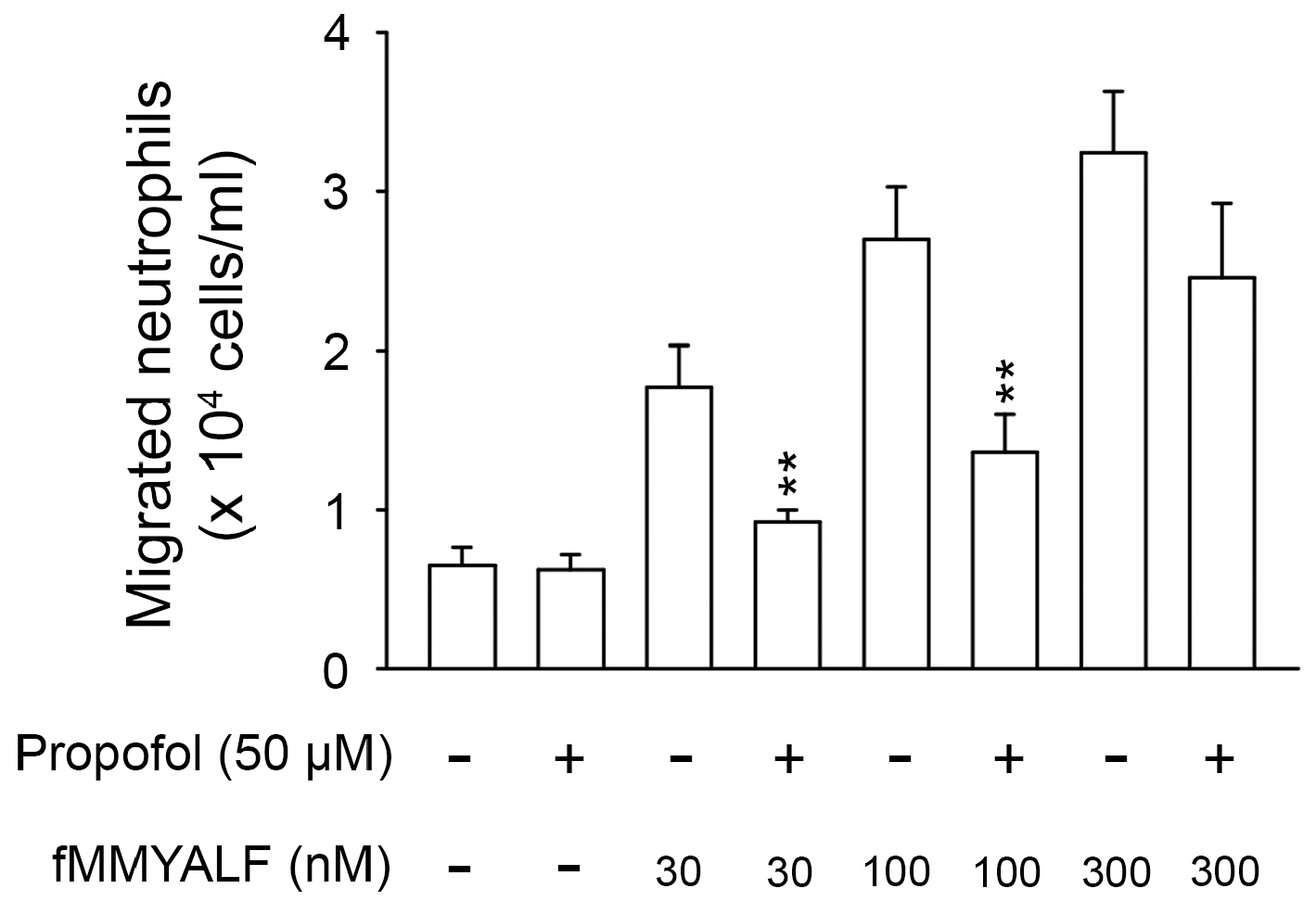
Effects of propofol on fMMYALF-induced neutrophil migration. fMMYALF-induced migrated neutrophils were counted (n = 5). Data information: Data are expressed as the mean ± SEM. **p < 0.01 (Student’s t-test).

### Propofol decreases intracellular Ca^2+^ mobilization and phosphorylation of protein kinase B and MAPKs

Notably, Ca^2+^ acts as an essential second messenger in neutrophil activations. Neutrophil stimulation with fMMYALF resulted in a prompt increase in the intracellular Ca^2+^ concentration ([Ca^2+^]_i_). Our results showed that propofol (50 μM) reduced the magnitude of the increase in [Ca^2+^]_i_ (Fig 6). The activation of phosphatidylinositol 3-kinase (PI3K) and subsequent triggering of protein kinase B (AKT) and mitogen-activated protein kinase (MAPK) pathways are involved in the FPR1 downstream signaling (Futosi et al, 2013). Propofol (50 μM) significantly reduced the phosphorylation of extracellular-signal-regulated kinase (ERK), p38, c-Jun N-terminal kinase (JNK), and AKT in human neutrophils treated with various concentrations of fMMYALF (Fig 7). Moreover, the reduction effect of propofol was reversed when the concentration of fMMYALF was increased.

**Figure 6.**
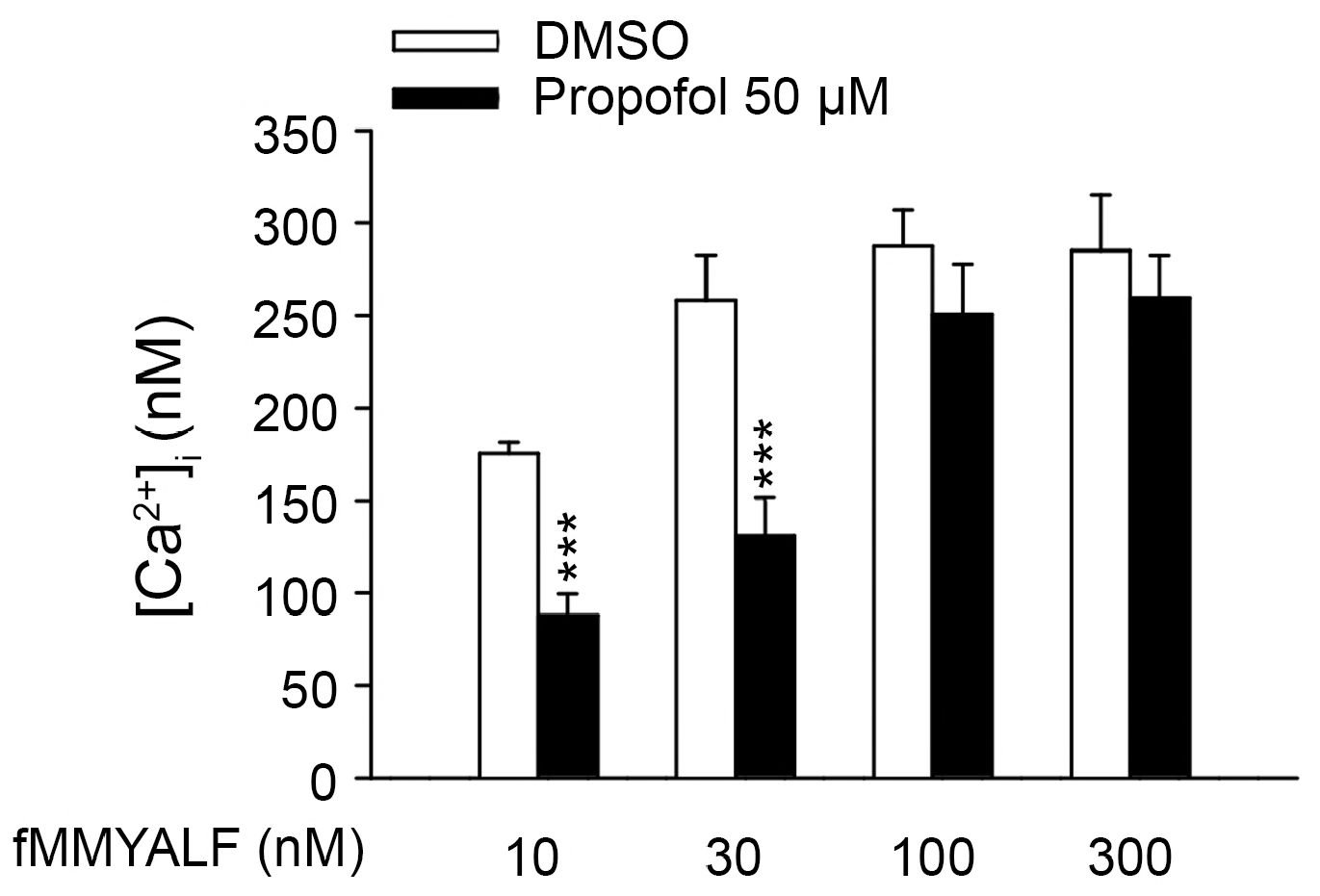
Effects of propofol on Ca2+ mobilization in fMMYALF-activated human neutrophils. The calcium concentrations in Fluo 3-loaded human neutrophils (n = 7). Data information: Data are expressed as the mean ± SEM. ***p< 0.001 (Student’s t-test).

**Figure 7.**
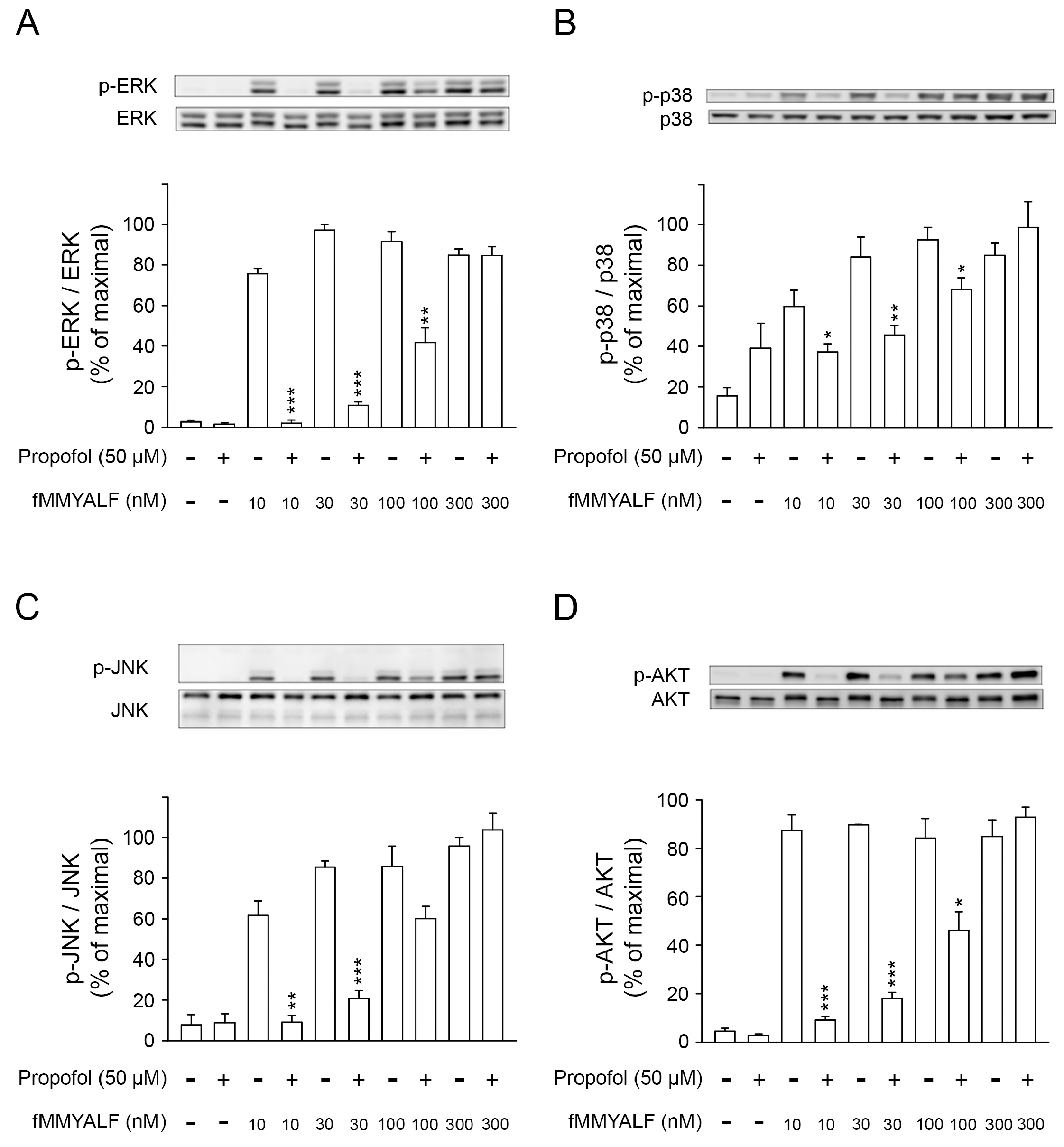
Effects of propofol on phosphorylation of AKT and MAPKs in fMMYALF-activated human neutrophils. A – D Western blotting of phospho- ERK, total ERK (A), phospho- p38, and total p38 (B), phospho-JNK, total JNK (C), phospho- AKT, total AKT (D). Quantification represents the phospho- protein/total protein ratios relative to the mean maximal ratio (n = 4). Data information: Data are expressed as the mean ± SEM. *p< 0.05, **p<0.01, ***p< 0.001 (Student’s t-test).

### Propofol attenuates sepsis-induced ALI and improves survival of septic mice

We used a sepsis-induced ALI murine model to evaluate whether propofol reduces the inflammation-associated lung impairment during sepsis. The lungs of mice with sepsis induced by intraperitoneally administered lipopolysaccharide (LPS) were collected and subjected to hematoxylin and eosin (HE) staining, immunohistochemical staining with myeloperoxidase (MPO) and Ly-6G, and an assay for MPO activity. The results of the lung injury assessment are presented for sham-operated mice treated with dimethylsulfoxide (DMSO; 50 μL, 10%, Group 1) or propofol (50 μL, 20 mg/Kg and 40 mg/Kg, Group 2), and mice with sepsis-induced ALI treated with DMSO (Group 3) or propofol (Group 4). As shown in Figure 8, sepsis resulted in alveolar wall disruption and edematous changes, which were alleviated by propofol. Sepsis also increased neutrophil infiltration, as evidenced by an increase in immunohistochemical MPO and Ly-6G expression. Intraperitoneal administration of propofol effectively reduced the increase in pulmonary neutrophil infiltration (Fig 8). Additionally, MPO activity, which is a quantitative indicator of neutrophil infiltration, exhibited a significantly increased in the lungs of septic mice. Administration of intraperitoneal propofol (20 and 40 mg/kg) significantly inhibited the rise in MPO activity in the lungs of septic mice (Fig 8A). Furthermore, a survival study was conducted, in which septic mice were challenged with 10 mg/kg LPS with or without propofol (50 μL; 10, 20 or 40 mg/kg). The survival rate was monitored for 7 days to evaluate overall survival (n = 6 for each group). Ultimately, within 3 days of the LPS challenge, all mice in control group perished; by contrast, 80% of mice treated with propofol (40 mg/kg) survived more than 7 days after the LPS challenge (Fig 9).

**Figure 8.**
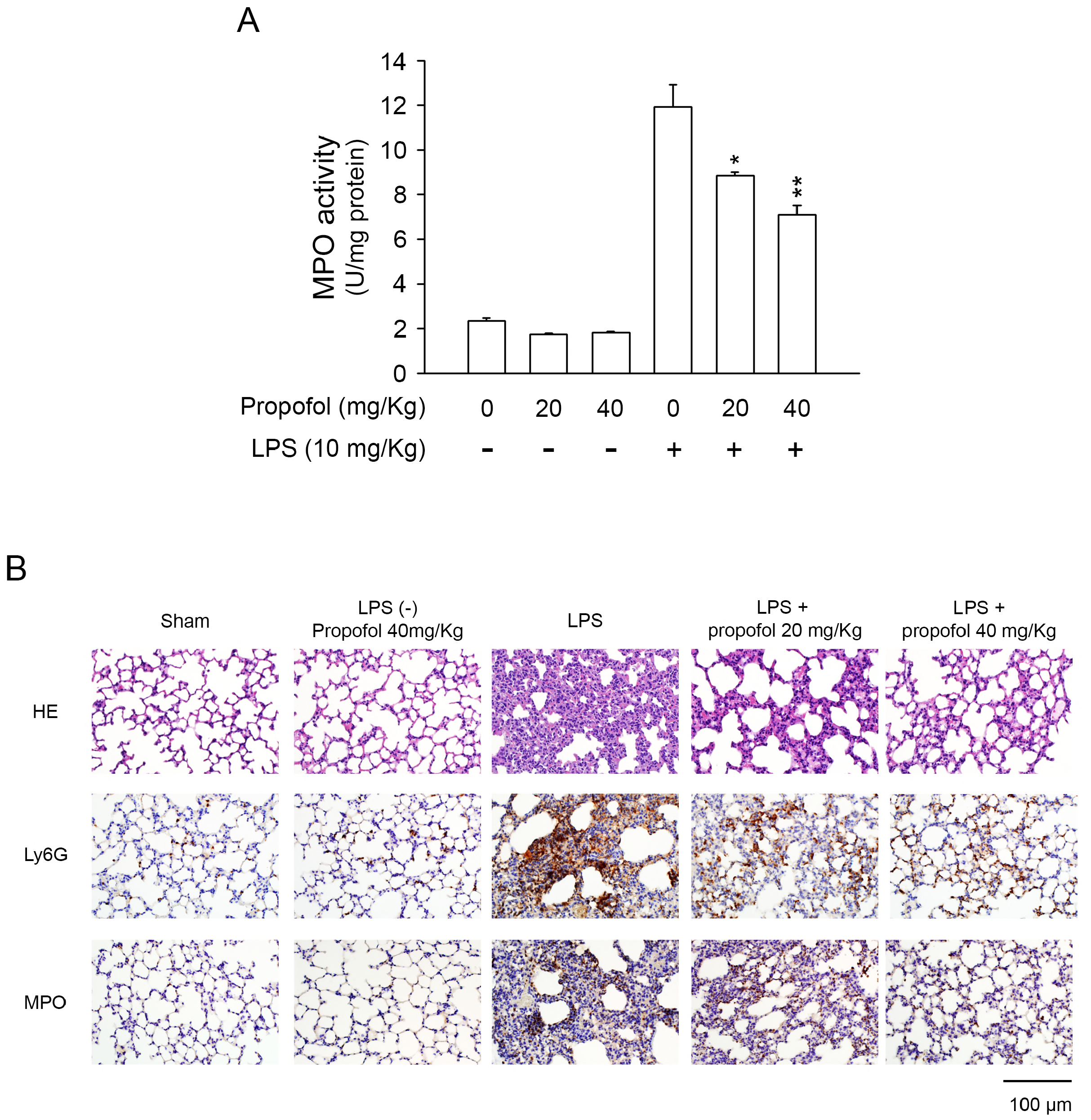
Propofol ameliorates neutrophil infiltration and lung damage in sepsis-induced acute lung injury (ALI).. A. MPO levels in the lungs of septic mice (n=7). B. Sections of lung tissue of hematoxylin and eosin (HE) staining (upper), immunohistochemical staining for neutrophilic Ly-6G proteins (middle) and MPO proteins (lower) (n =6 mice for each group). Scale bars: 100 μm. Data information: Data are expressed as the mean ± SEM. *p< 0.05, **p<0.01 (Student’s t-test).

**Figure 9.**
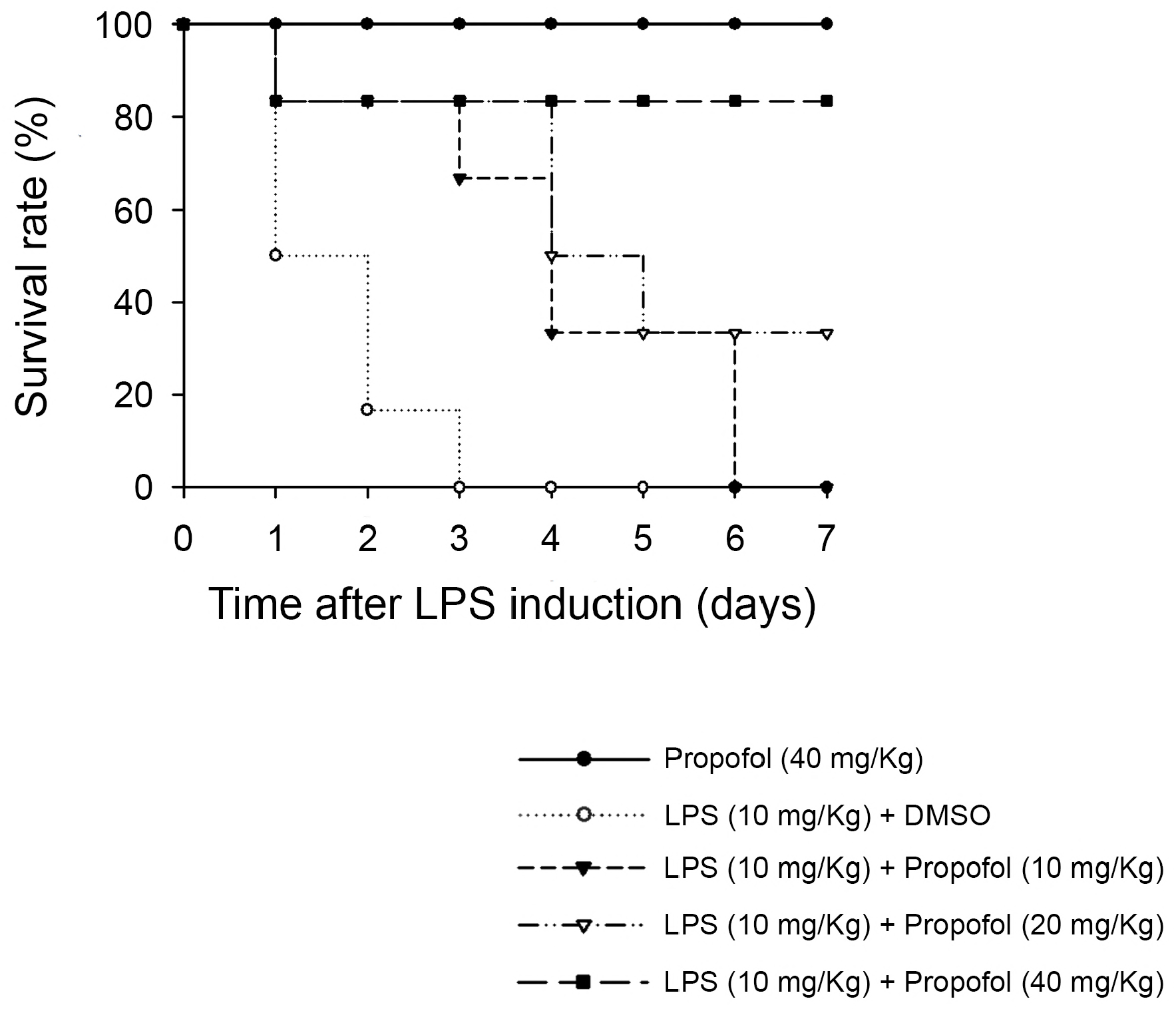
Propofol ameliorates LPS-induced mortality in mice. The survival rate of mice (n = 6 for each group). 100% of the mice died within 3 days after the LPS challenge, whereas 80% of mice treated with propofol survived for 7 days after the LPS challenge.

## Discussion

Propofol is commonly used in operation rooms or intensive care units (ICUs) for critically ill patients. Various clinical trials have demonstrated that the antioxidant, anti-inflammatory, and free-radical-scavenging properties of propofol can provide clinical benefits. For examples, the use of propofol has been reported to reduce the incidence of postoperative cognitive dysfunction, maintain an appropriate hemodynamic status, shorten ICU stays, and increase the lung compliance of patients who underwent major surgery or cardio-pulmonary bypass (An et al, 2008; Huang et al, 2011; Landis et al, 2014; Qiao et al, 2015; Sayed et al, 2015). Nevertheless, the detailed molecular mechanism that governs these protective effects of propofol is still incompletely understood. The present study illustrated that propofol competitively inhibits the interaction between FPR1 and fMMYALF, which is a mitochondria-derived DAMP released from damaged cells (Eleftheriadis et al, 2016; Krysko et al, 2011; Raoof et al, 2010). Therefore, propofol can reduce degranulation, respiratory burst, and cell migration, which are all inflammatory responses of neutrophils that can be observed in critically-ill patients caused by traumatic or surgical-associated injury.

The human immune is a complex system whose primary aim is to protect human the body from invading pathogens. To achieve this aim, recognition of pathogenic microbes is the first and most critical step, followed by the initiation of a series of responses required for eliminating the invading pathogens. Nevertheless, while invading pathogens that cause tissue injury must be eliminated, the commensal microorganisms that are required for host survival should be tolerated. In addition, the immune system plays a crucial role in maintaining hemostatic tissue function. Sterile tissue injury arising from traumatic or surgery-associated injury must be detected and repaired. In other words, with the emergence of the “danger model” in the scientific community, researchers have come to understand that the immune system depends on various pattern-recognition receptors (PRPs) to distinguish danger and nondanger patterns, rather than self and nonself patterns (Matzinger, 1994). The common causes of critical illness, including sepsis, trauma, surgery-related cell damage, hypoxia, and ischemia, also cause uncontrolled cell or tissue damage. Consequently, intracellular molecules, some of which are considered as DAMPs, are either actively or passively discharged into the surrounding tissue and circulation (Matzinger, 1994; Oppenheim & Yang, 2005). Moreover, because critically ill patients exhibit immune-associated complications, DAMPs have crucial implications in the prognosis of these patients and are considered possible therapeutic targets for anti-inflammatory compounds.

In the past decade, studies have demonstrated that the mitochondrial molecules released into circulation act as DAMPs; therefore, they contribute to DAMP-mediated immune stimulation (Krysko et al, 2011; Raoof et al, 2010). Mitochondria originated from engulfed prokaryotic cells (Gray et al, 1999; Zimmer, 2009). Mitochondrial protein synthesis is similar to that of their prokaryotic ancestor and begins with the H-formyl methionine residue (Marcker, 1965; Raoof et al, 2010), which is absent in the cytosol. The fMMYALF is an *N*-formyl peptide that corresponds to the *N* terminus of mitochondrial NADH dehydrogenase subunit 6. Prior studies have reported that fMMYALF, similar to fMLF of prokaryotic origin, is a portent activator that stimulates immune effector cells via FPRs (Crouser et al, 2009; Rabiet et al, 2005; Raoof et al, 2010). It induces p38, p44, and p42 MAPK phosphorylation, resulting in the release of proinflammatory signals, including matrix metalloproteinase-8 and interleukin 8 (Raoof et al, 2010; Zhang et al, 2010). In the present study, we used the fMMYALF for human neutrophil stimulation in order to investigate the immunomodulatory effects of propofol on the critically ill patients with inflammatory responses.

The receptor binding assay used in our previous study demonstrated that propofol competitively binds to FPR1, thus blocking the downstream signal transduction of FPR1 and inhibiting neutrophil immune activities (Yang et al, 2013). In this study, we assessed the pharmacological action of fMMYALF in the presence of propofol. Cellular inflammatory responses, including elastase release, superoxide release, ROS production, and cell migration were assessed at different concentrations of the stimulant (fMMYALF) and constant concentration of the inhibitor (propofol). The addition of propofol (50 μM) causes a parallel rightward shift of the fMMYALF response curve (Fig 1B, 2B), which represents an increase in the EC50 of the agonist (fMMYALF). In other words, propofol reduces the potency but not the maximal response of fMMYALF.

FPRs is a member of Gi/o protein-coupled receptor. Ligand interaction with FPRs initiates the dissociation of the Gα subunit from the Gβγ subunit. Intracellular Ca^2+^ mobilization occurs downstream of G-protein activation and is crucial to respiratory burst in human neutrophils (Berridge et al, 2000; Brechard & Tschirhart, 2008). Our data demonstrated that propofol reduced the fMMYALF-induced mobilization of intracellular Ca^2+^. The activation of PI3K and the subsequent production of phosphatidylinositol (3,4,5) P3 (PIP3) and AKT are other major signal transduction event associated with FPRs and mediated by Gβγ dimers (Stephens et al, 1994). Furthermore, the p38 and ERK MAPKs are downstream mediators of the FPR signaling pathway (Heit et al, 2008; Mocsai et al, 2000). They are strongly activated on neutrophil stimulation with FPR1 agonists. The p38 promotes the chemotactic migration of neutrophils (Hannigan et al, 2001). Western blot analysis in our experiments revealed that propofol reduced fMMYALF-induced phosphorylation of ERK, p38, JNK, and AKT. Taken together, the preceding evidence demonstrates that propofol can effectively reduce fMMYALF-induced human neutrophil inflammatory responses by directly interfering with the binding of fMMYALF to FPR1, thereby blocking the subsequent signal transduction.

Despite considerable progress in elucidating the mechanisms of sepsis and sepsis-associated ALI as well as the development of novel intervention therapies in ICUs, sepsis and sepsis-induced ALI remain leading causes of mortality in critically ill patients (Matthay et al, 2012). Recruitment and infiltration of neutrophils is considered as a critical process in ALI (Grommes & Soehnlein, 2011; Zemans & Matthay, 2017). Because FPR1 plays a vital role in the process of neutrophil migration during acute and chronic inflammation (Tsai et al, 2016), researchers have carefully investigated the importance of FPR1 in neutrophil recruitment during the development of LPS-induced ALI (Wenceslau et al, 2016). Specifically, a prior study demonstrated that the lung infiltration of neutrophils was markedly reduced in the FPR1^−/−^ mice (Grommes et al, 2014). Moreover, we and other researchers have shown that cyclosporine H, an FPR1 antagonist, can pharmacologically reduce the neutrophils recruitment, thus affording protection effect against LPS-induced ALI (Grommes et al, 2014; Yang et al, 2017). In the present study, propofol was shown to reduce pulmonary edema, neutrophil infiltration, and MPO activity in our experimental murine model of LPS-induced ALI. Additionally, propofol reduces the mortality rate of mice with endotoxemic ALI. Taken together, this evidence shows that FPR1 is an important target of anti-inflammatory strategies and provides protection against LPS-induced ALI.

We concluded that the inhibitory effects of propofol on fMMYALF-induced neutrophil activation are mediated by competition with FPR1, which inhibits the downstream signaling of receptors (Fig 10). Our experiments performed using a septic mouse model revealed that propofol, an FPR1 antagonist, may be beneficial for treating patients who are critically ill with sepsis and sepsis-associated ALI.

**Figure 10.**
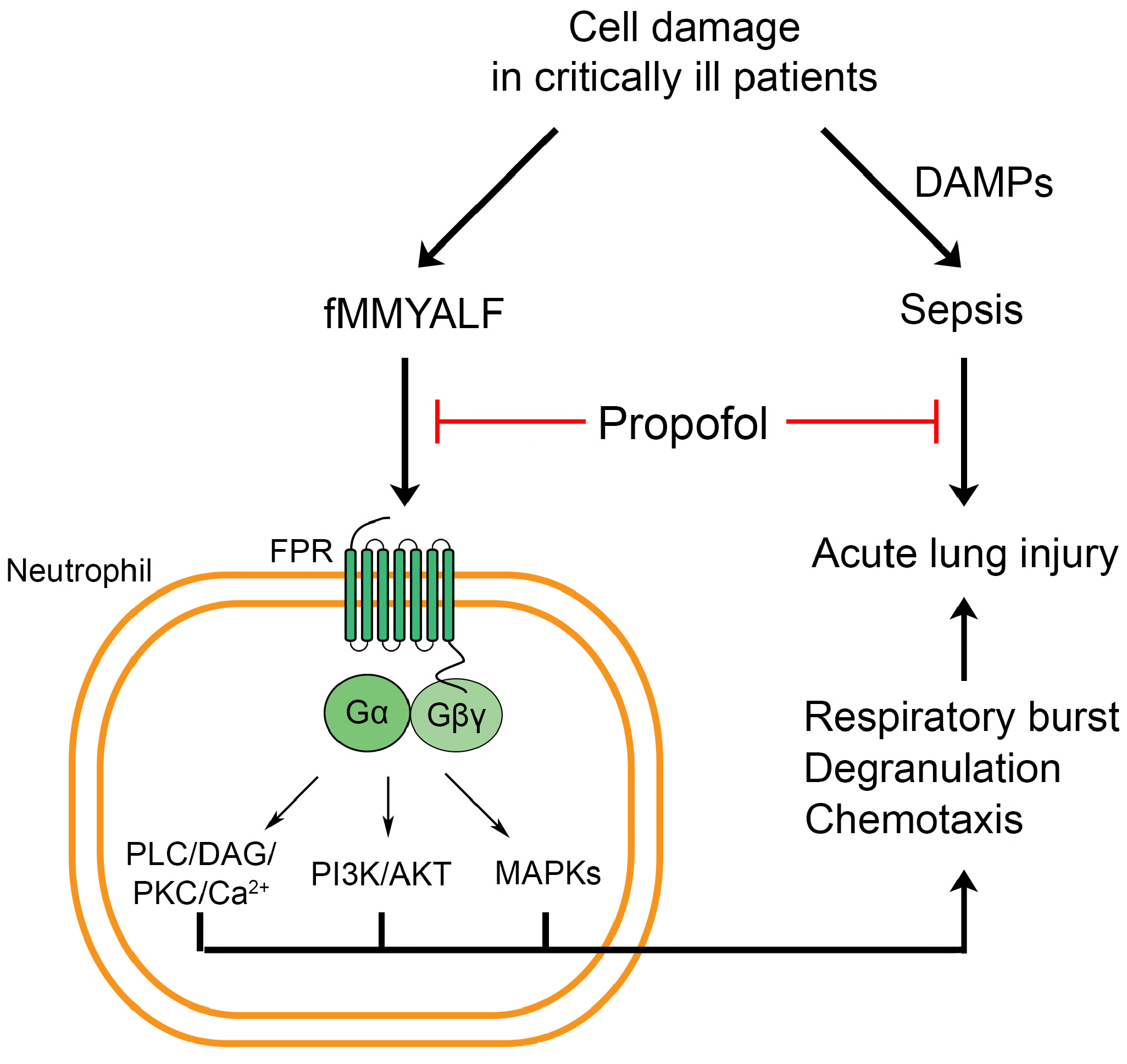
Schematic diagram illustrating the therapeutic effects of propofol in human neutrophils and sepsis-induced acute lung injury (ALI). Propofol inhibits respiratory burst, degranulation, and chemotactic migration in fMMYALF-activated human neutrophils by interfering with the binding of fMMYALF to FPR1, which inhibits the downstream signaling of receptors. Propofol attenuates sepsis-induced ALI and mortality in mice.

## Materials and methods

### Reagents

Propofol (2,6-diisopropylphenol) was obtained from Sigma–Aldrich (St. Louis, MO, USA). Dihydrorhodamine 123 (DHR 123) and fluo-3 acetomethoxyester (fluo-3/AM) were purchased from Molecular Probes (Eugene, OR). Antibodies directed against phospho-ERK, ERK, phosphor-AKT (ser-473), and AKT were purchased from Cell Signaling (Beverly, MA, USA). An anti-p38 antibody was obtained from Santa Cruz Biotechnology (Santa Cruz, CA, USA). All other pharmacologic agents were purchased from Sigma–Aldrich.

### Preparation of human neutrophils

The research protocol was granted approval by the institutional review board of Chang Gung Memorial Hospital. After obtaining written informed consent, human neutrophils were isolated from heparinized venous blood donated by healthy participants aged 20 to 30 years who had reported no systemic disease within 1 week before blood collection. The procedures used for human neutrophil isolation have been described previously (Yang et al, 2013). Briefly, the blood samples were subjected to dextran sedimentation and Ficoll–Hypaque centrifugation, followed by hypotonic lysis of contaminating red blood cells. The segregated neutrophils were suspended and stored in pH 7.4 Ca^2+^-free Hank’s balanced salt solution (HBSS) at 4°C prior to the experiments. We employed the Wright–Giemsa stain to confirm the purity of neutrophil suspension. Trypan blue exclusion was applied, and a viability of >98% was established.

### Assessment of the elastase release

We measured elastase release to determine the degranulation function of activated neutrophils (Hwang et al, 2006). The elastase substrate was methoxysuccinyl-Ala-Ala-Pro-Val-p-nitroanilide. Human neutrophils (6 × 10^5^ cells/mL) were mixed with the substrate (0.1 mM) in HBSS with CaCl_2_ (1 mM) at 37°C. The neutrophils were treated with DMSO or propofol (5–100 μM) for 5 min. They were then activated by different concentrations of fMMYALF (30–30000 nM) in pretreatment with cytochalasin B (0.5 μg/mL). A spectrometer (Hitachi U-3010, Tokyo, Japan) was employed to continuously record the change in absorbance at 405 nm.

### Determination of superoxide release

We used ferricytochrome c, which can be reduced by superoxide, to evaluate the superoxide release in human neutrophils (Chen et al, 2014). Briefly, neutrophils (6 × 10^5^ cells/mL) were incubated in HBSS containing ferricytochrome *c* (0.5 mg/mL) and CaCl_2_ (1 mM) at 37°C. Afterward, they were treated with DMSO or propofol (5–100 μM) for 5 min. Neutrophils activation was achieved by adding fMMYALF (0–30000 nM) with pretreatment of cytochalasin B (1 μg/mL). We continuously monitored the changes in absorbance at 550 nm resulting from a ferricytochrome *c* reduction with a double beam spectrophotometer (Hitachi U-3010) with stirring. We calculated the superoxide release as the difference between the reaction with superoxide dismutase (100 U/mL) and that without superoxide dismutase divided by the extinction coefficient for ferricytochrome *c*’s reduced form (Chen et al, 2014).

### Assay of intracellular ROS production

Nonfluorescent DHR 123 can be converted to fluorescent rhodamine 123 by ROS; therefore, we determined the intracellular ROS concentration by measuring the fluorescence of rhodamine 123. We mixed human neutrophils (6 × 10^6^ cells/mL) with DHR 123 (2 μM) at 37 °C for 15 min. They were subsequently treated with DMSO or propofol (50 μM) for 5 min. Activation of neutrophils was performed by adding fMMYALF (30–3000 nM) for another 5 min. Subsequently, the reaction was terminated by putting the samples on ice. Flow cytometry was applied to ascertain the change in fluorescence.

### Measurement of total ROS release

We used the luminol-enhanced chemiluminescence method to measure the total ROS concentration. We incubated human neutrophils (2 × 10^6^ cells/mL) with luminol (37.5 μM) and horseradish peroxidase (6 U/mL) for 5 min at 37°C. Subsequently, they were treated with DMSO or propofol (50 μM) for 5 min. Activation of neutrophils was executed using fMMYALF (30–3000 nM). Chemiluminescence was detected using a Tecan Infinite F200 Pro 96-well chemiluminometer (Tecan Group, Männedorf, Switzerland).

### Determination of LDH release

In order to exclude the possible cytotoxic effect of the combination of propofol and fMMYALF, LDH activity was assessed with a commercial LDH assay kit (Promega, Madison, WI, USA). Briefly, human neutrophils (1.2 × 10^6^ cells/mL) were treated with propofol (50 μM) for 5 min and fMMYALF (10–3000 nM) for another 10 min. The LDH assay reagents were then added to the supernatant and incubated for another 30 min. The fluorescence signal was assessed and compared with the total LDH activity, which was defined by measuring the fluorescence of lysed neutrophils treated with lysing solution (0.1% Triton X-100) for 30 min.

### Evaluation of cell migration

The chemotaxis of human neutrophils was evaluated using a microchemotaxis chamber with a 3-μm filter (Millipore, Darmstadt, Germany). Neutrophils (2.5 × 10^6^ cells/mL) were treated with DMSO or propofol (50 μM) for 5 min at 37°C in the upper chamber. The lower chamber contained various concentrations of fMMYALF (30–300 nM). After incubation in 5% CO_2_ for 120 min, a cell counter (Moxi, Orflo, Ketchum, ID, USA) was used to ascertain the number of migrating neutrophils.

### Determination of intracellular calcium concentration

We cocultured neutrophils (3 × 10^6^ cells/mL) with fluo-3/AM (2 μM) for 45 min at 37°C. The neutrophils were then centrifuged and resuspended in HBSS with CaCl_2_ (1 mM). We then treated these labeled cells with DMSO or propofol (50 μM) for 5 min. The cytosolic calcium levels in response to fMMYALF (10–300 nM) were obtained using a spectrophotometer (Hitachi F-4500). The excitation and emission wavelengths were 488 and 520 nm, respectively. In each experiment, 20 mM ethylene glycol tetraacetic acid and 0.05% Triton X-100 were added to obtain the minimum and maximum fluorescence values, respectively.

### Immunoblotting analysis

Human neutrophils were treated with DMSO or propofol (50 μM) for 5 min and subsequently stimulated with fMMYALF (10–300 nM) for 30 s, followed by placing the cells on ice to stop the reaction. Cells were then centrifuged at 4°C, and the precipitated pellet was lysed in lysis buffer (150 μL) containing NaCl (100 mM), EDTA (1 mM), HEPES (50 mM, pH 7.4), Na_3_VO_4_ (2 mM), p-nitrophenyl phosphate (10 mM), 2-ME (5%), PMSF (1 mM), protease inhibitor cocktails (1%, Sigma–Aldrich), and Triton X-100 (1%). Following sonication, the lysates underwent centrifugation (14,000 rpm at 4°C for 20 min) and were subjected to sodium dodecyl sulfate–polyacrylamide gels (12%) to separate the proteins. The proteins separated through electrophoresis were transferred onto nitrocellulose membranes, followed by blocking the membrane with 5% nonfat milk in a mixture of Tris-buffered saline and Tween 20. The membranes were incubated in solution containing relevant primary and horseradish peroxidase-conjugated secondary antibodies (Cell Signaling Technology, Beverly, MA). An enhanced chemiluminescence solution (Amersham Biosciences) was added to the membranes, and proteins were detected and quantified using the BioSpectrum Imaging System (UVP, Upland, CA, USA).

### LPS-induced sepsis model

All animal experiments were approved by the Institutional Animal Care and Use Committee of Chang Gung University. BALB/c mice (male, 20–25 g, 7–8 weeks old) were acquired from BioLASCO (Taipei, Taiwan). They were housed in an air-conditioned room under a 12-h light–dark cycle. They were provided with 1 week to adapt to the housing environment prior to the experiment. Mice were randomly divided into four groups as follows: Group 1, sham-operated mice treated with DMSO (50 μL, 10%); Group 2, sham-operated mice treated with propofol (50 μL, 20 mg/Kg and 40 mg/Kg); Group 3, septic mice treated with DMSO (50 μL, 10%); and Group 4 (50 μL, 20 mg/Kg and 40 mg/Kg), septic mice treated with propofol. Propofol was dissolved in DMSO and then diluted 10-fold using normal saline. Three doses of DMSO or propofol were administered to the mice intraperitoneally every 30 min. Following the second administration of DMSO or propofol, the mice were challenged with a single 200-μL dose of LPS (10 mg/Kg, *Escherichia coli*, serotype O111:B4, Sigma–Aldrich, USA) or 0.9% saline (sham operation) administered intraperitoneally. The mice were humanely sacrificed under isoflurane inhalation–administered anesthesia 20 h after LPS administration. Two segments of the right upper lobe of the lung were fixed in 10% neutral buffered formalin. They were subsequently set in paraffin blocks, and then cut into 5-μm-thick sections for histological and immunohistochemical analysis. The other tissues were frozen at −80°C until the assay. In survival study, septic mice were challenged with a single 200-μL dose of LPS (10 mg/kg) with or without three doses of 50 μL propofol (10, 20, or 40 mg/kg). The survival rate was monitored for 7 days (n = 6).

### MPO activity assay

To measure the MPO activity, lung tissues were homogenized in phosphate buffered saline containing 0.5% hexadecyltrimethylammonium bromide (Sigma–Aldrich, USA) by using the MagNA Lyser Instrument (Roche, Germany). After being centrifuged (12,000 ×*g*) at 4°C for 20 min, the supernatant fluids were incubated with 0.2 mg/mL *o*-dianisidine dihydrochloride containing 0.001% H_2_O_2_. The absorption was spectrophotometrically determined at 405 nm. The MPO activity was normalized to the protein concentration. The total protein concentration in the homogenate was ascertained by employing the Bradford method (Bio-Rad, USA).

### Histology and Immunohistochemistry

The paraffin-embedded sections were deparaffinized and stained using HE. For immunohistochemical staining, the sections were incubated with a Ly6G antibody (eBiosciences, USA) and MPO antibody (Abcam, UK). Secondary labeling was achieved using the SuperPicture Polymer Detection kit (Thermo Fisher, USA). We observed the sections by using an Olympus BX51 microscope. Additionally, a DP12 digital camera was used to record images.

### Statistical analysis

Data are presented from four to seven samples used for each specific experiment. The data are expressed as means ± SEM. Statistical analysis was executed using Student’s *t* test. The SigmaPlot (Systat Software, San José, CA, USA) software was used for all computation. A *p*-value of 0.05 was considered statistically significant.

## Acknowledgments

This research was financial supported by the grants from the Ministry of Science Technology (MOST 106-2320-B-255-003-MY3 and MOST 104-2320-B-255-004-MY3), Ministry of Education (EMRPD1G0231), and Chang Gung Memorial Hospital (CMRPF1F0011∼3, CMRPF1F0061∼3, CMRPF1G0241∼3, and BMRP450), Taiwan. The funders had no role in study design, data collection and analysis, decision to publish, or preparation of the manuscript.

## Author contribution

CY Chen, YF Tsai, and WJ Huang carried out most *in vitro* assays. CY Chen, YF Tsai, and SH Chang performed most animal studies. CY Chen and TL Hwang were responsible for the overall design, analyzed data, and wrote the article. All authors analyzed the results and approved the final version of the manuscript.

## The Paper Explained

Problem: Sedative agents should be selected carefully for anesthetizing critically ill patients who are at high risk of secondary infection and sepsis. Propofol has anti-inflammatory effects that benefit critically ill patients who require anesthesia. However, the mechanism and therapeutic effect of propofol on neutrophil-dominant acute lung injury remain incompletely understood.

Results: Here we found that propofol competitively inhibited various neutrophil activities by blocking the interaction between endogenous N-formyl peptide and its receptor, FPR1, thus inhibiting downstream signaling. Propofol also alleviated the severity of sepsis-associated acute lung injury in mice and improved the survival.

Impact: Our *in vitro* and septic mouse model experiments revealed that the anesthetic agent propofol, an FPR1 antagonist, may be beneficial for treating patients who are critically ill with sepsis and sepsis-associated ALI.

